## Supplementary figures and images for "Enhanced RNA replication and pathogenesis in recent SARS-CoV-2 variants harboring the L260F mutation in NSP6"

### Figure S1

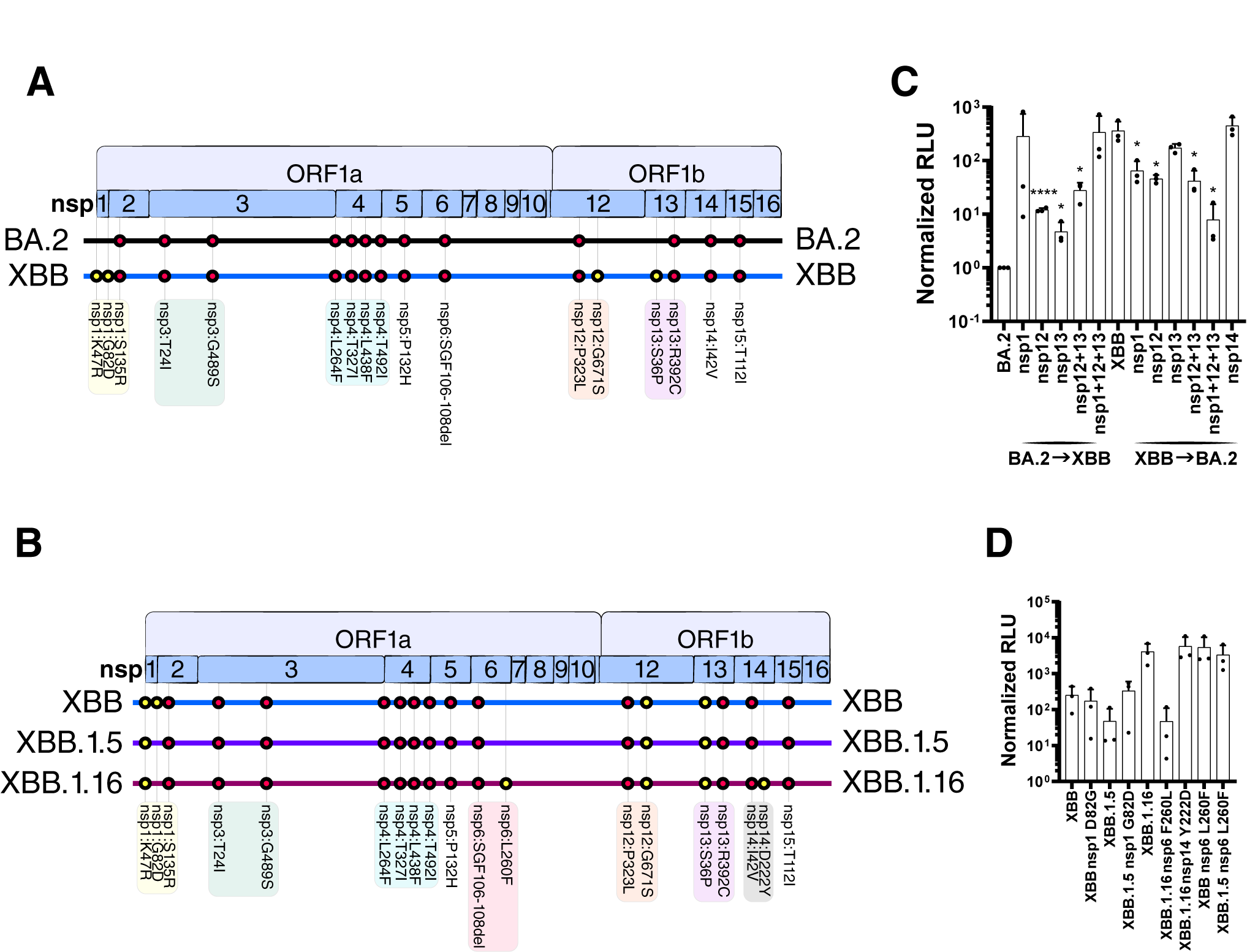

### Figure S2

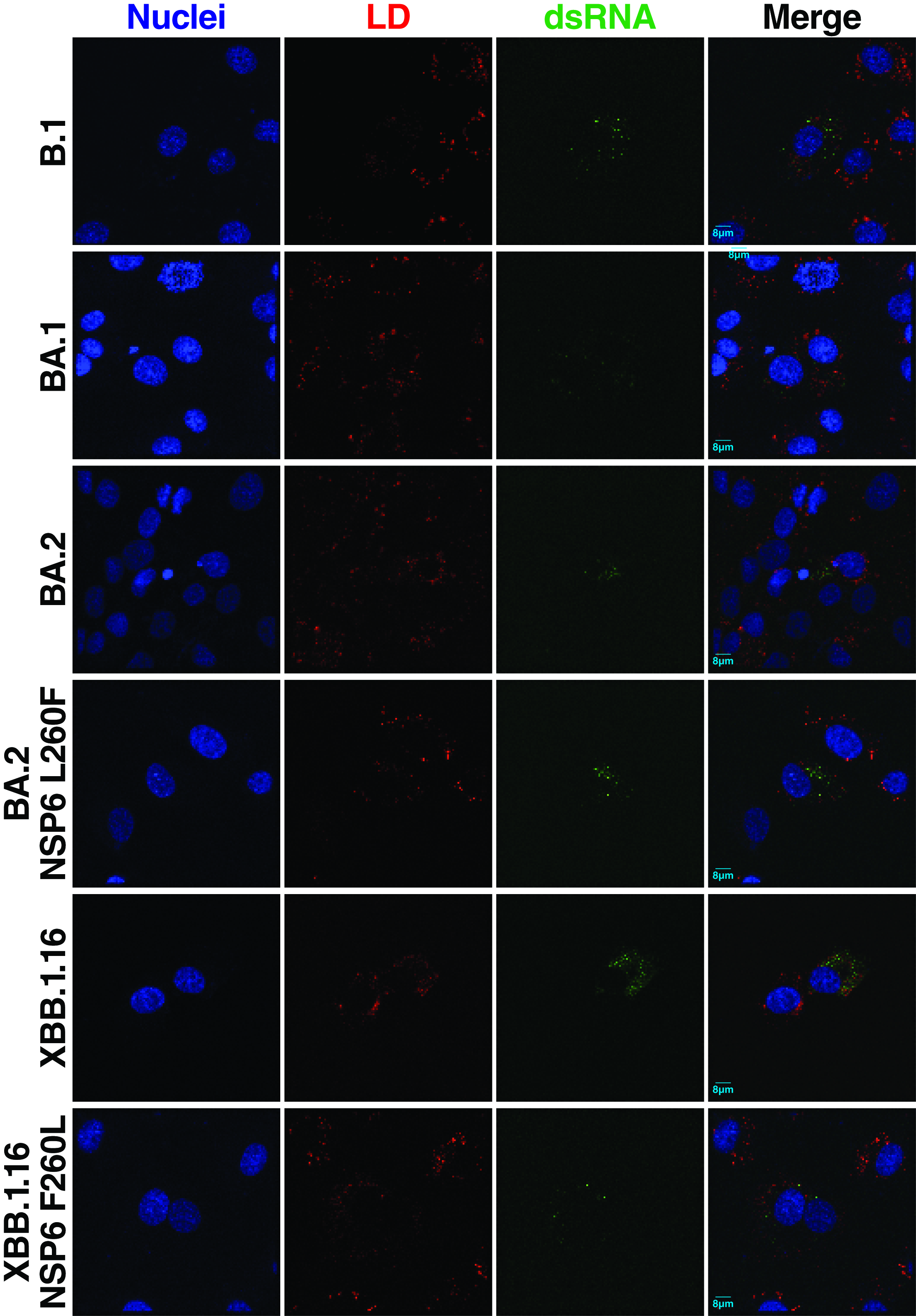

### Figure S3

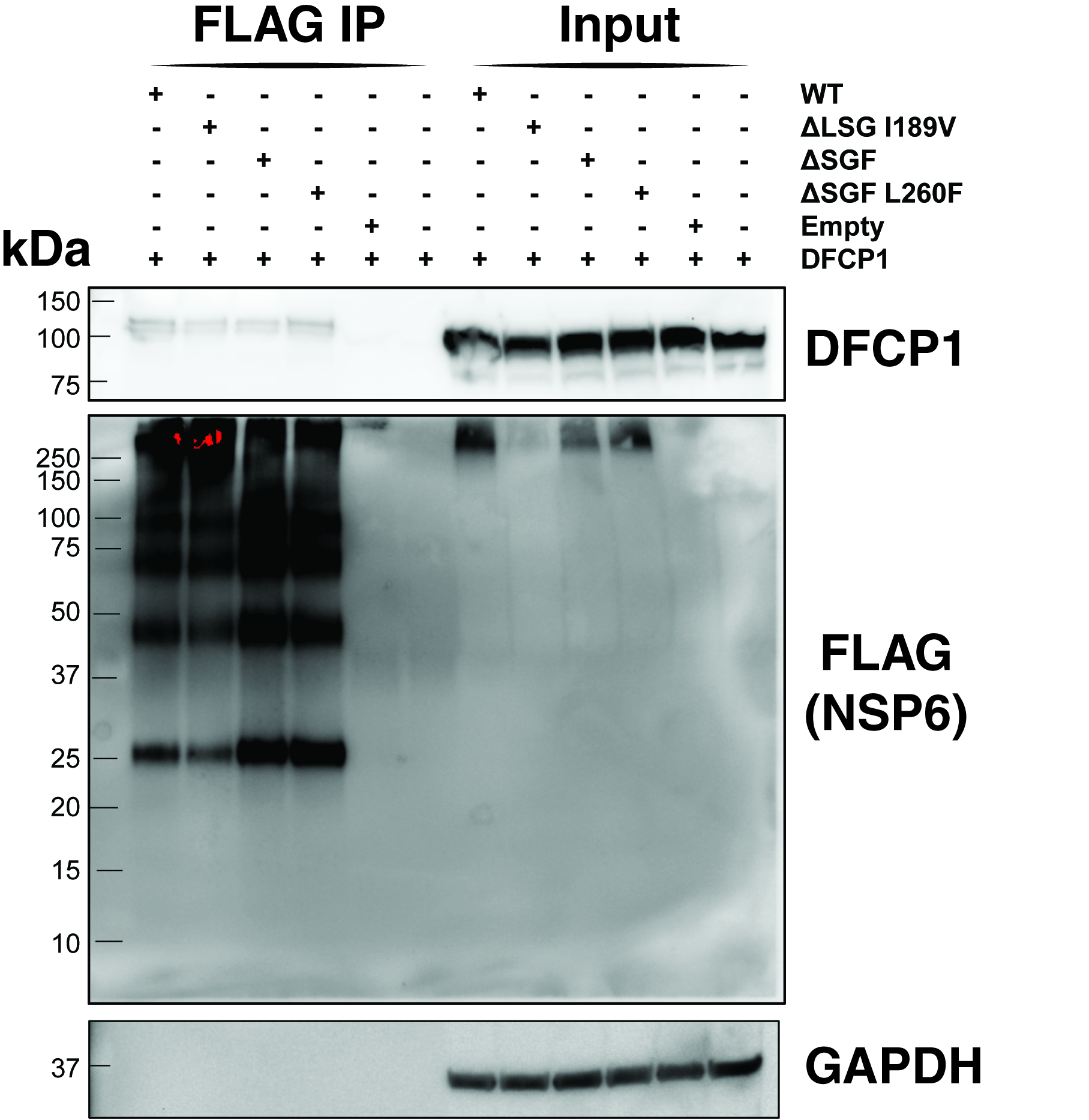

### Figure S4

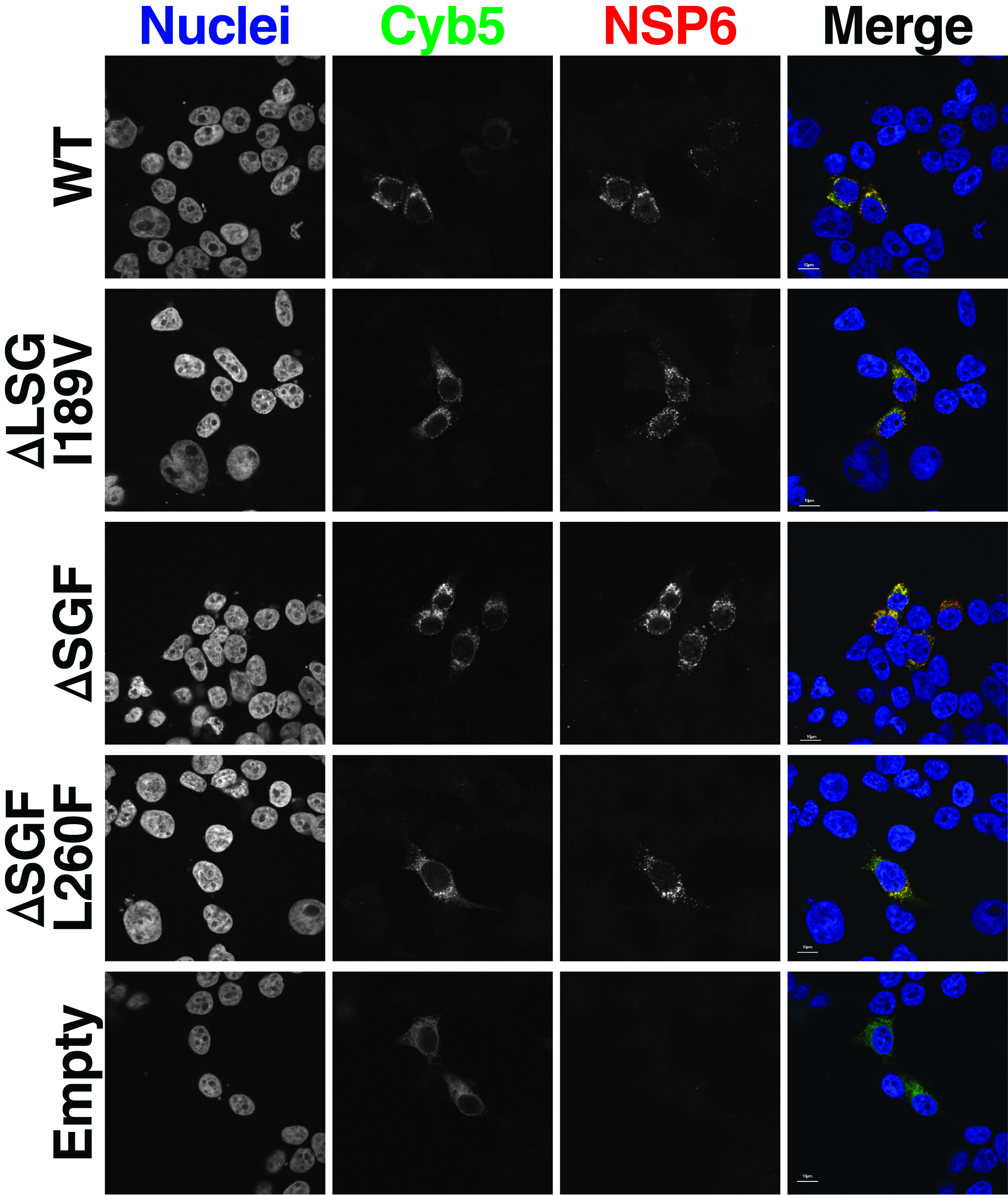

### Figure S5

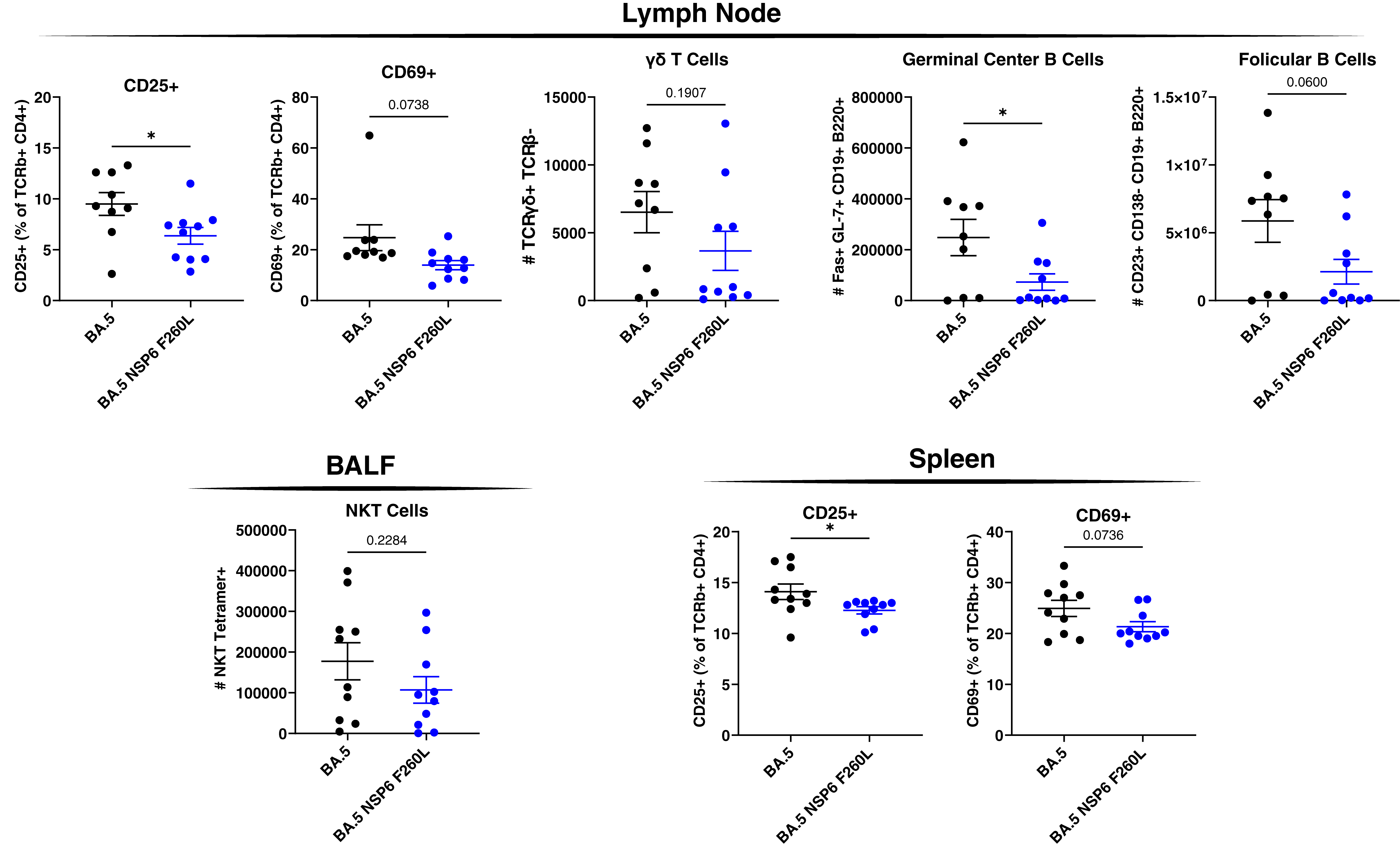

### Figure S6

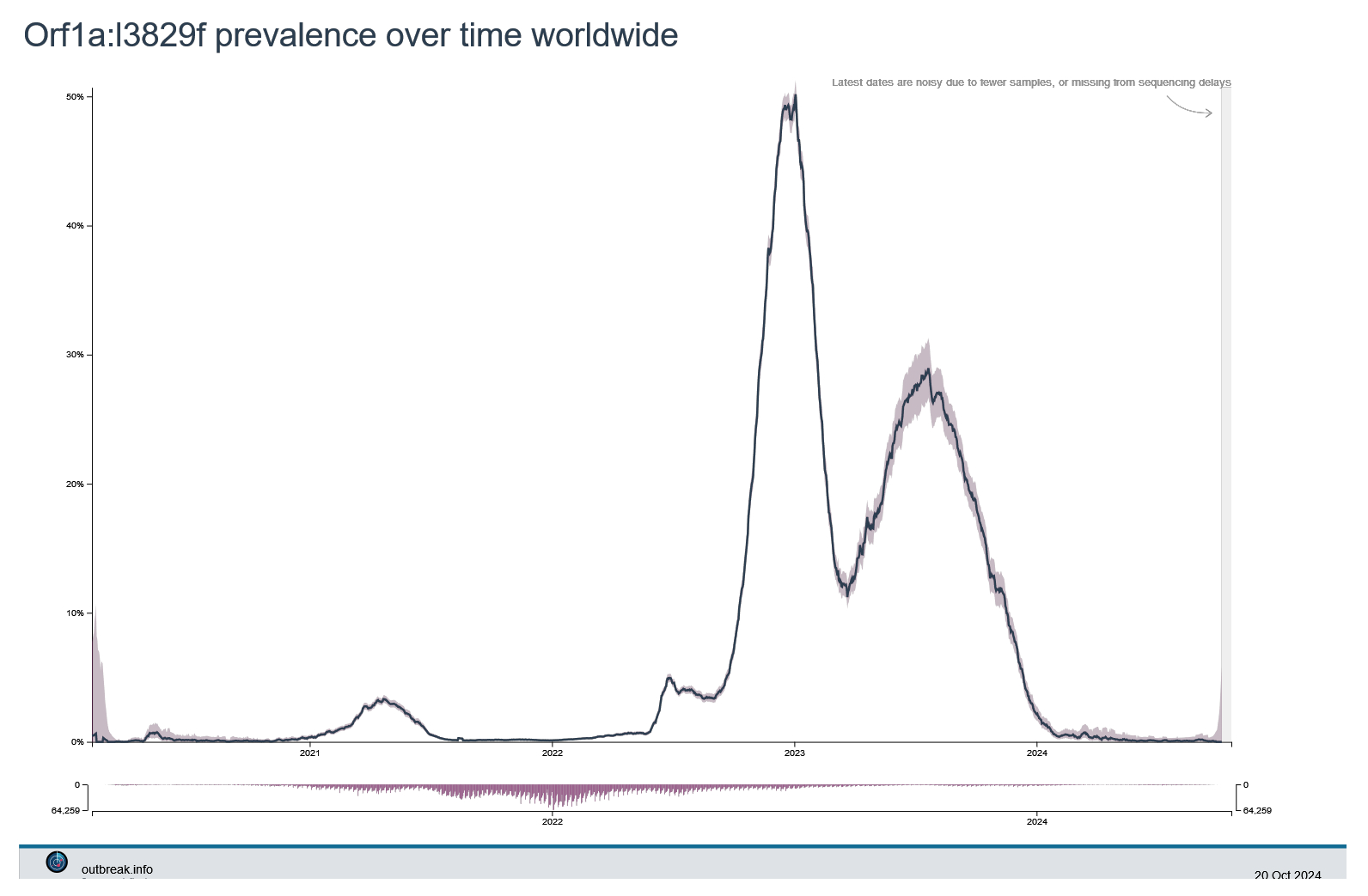
